# Existence of log-phase Escherichia coli persisters and lasting memory of a starvation pulse

**DOI:** 10.1101/2020.09.17.301598

**Authors:** Mikkel Skjoldan Svenningsen, Sine Lo Svenningsen, Michael Askvad Sørensen, Namiko Mitarai

## Abstract

The vast majority of a bacterial population is quickly killed when treated with a lethal concentration of antibiotics. The time scale of this killing is often comparable with the bacterial generation time before addition of antibiotics. Yet, a small subpopulation typically survives for an extended period. However, the long-term killing dynamics of bacterial cells has not been fully quantified even in well-controlled laboratory conditions. We constructed a week-long killing assay and followed the survival fraction of *Escherichia coli* K12 exposed to a high concentration of ciprofloxacin. We found that long-term survivors were formed during exponential growth, with some cells surviving at least 7 days. The long-term dynamics contained at least three timescales, which greatly enhances predictions of the population survival time compared to the biphasic extrapolation from the short term behavior. Furthermore, we observed a surprisingly long memory effect of a brief carbon starvation pulse, which was dependent on the (p)ppGpp synthase *relA*. Specifically, one hour of carbon starvation prior to antibiotics exposure increased the surviving fraction by nearly 100-fold even after 4 days of ciprofloxacin treatment.

## 1. Introduction

Bacterial populations are quickly decimated during a typical antibiotics assault. Within a few generation times, the far majority of cells will be dead. However, it is typically recommended to use extended durations of treatment, ranging from several days to months, prolonging the exposure time of bacterial pathogens to the antibiotic [1]. The WHO is now considering the benefits of shortening the duration of antibiotics administration while still keeping the treatment effective, due to concerns of increasing antibiotic resistance occurring as a consequence of increased exposure [1]. To find the optimal treatment duration, one needs to understand the killing dynamics of bacteria when exposed to antibiotics, especially the bacterial cells surviving for longer times. The long-term survivors are typically referred to as persister cells, a subgroup of cells that survive antibiotics for an extended period compared to the average of the population, but have not acquired mutations that make them resistant to the antibiotic [2, 3]. Most research on persister cells is done within the well-defined conditions of the laboratory, but despite these strongly simplified conditions, and more than seventy years of research, laboratory persisters are still far from understood [4, 5].

One pending question is whether and how much persisters form spontaneously during the exponential growth phase. Such persisters are called type-II [3] or spontaneous [4] persisters. It was repeatedly shown that stress-triggered (or type I) persisters are formed in high numbers during the stationary phase, but research on spontaneous persister formation during the exponential phase is sparse [4]. The research has mostly been confounded by a lack of careful attention to the presence of stationary phase cells carried over from the starter cultures, which artificially elevated the persister fraction of exponential cultures [5, 6, 4]. One carefully executed study showed that no *E. coli* persister cells were formed during fast exponential growth in rich medium [7], whereas other studies merely showed reduced levels during exponential growth [3, 5]. A benefit of analysing the exponential growth phase is the well-defined physiology of this state [8, 9]. This makes it possible to vary the growth physiology in a controlled manner, especially by varying the growth rate through culturing bacteria in media of different nutrient quality. It was previously shown that the bacterial growth rate strongly correlates with the death rate during the initial period of killing with beta-lactams [10, 11, 12]. This poses the additional questions of how the growth rate at the time of antibiotics exposure affects the shortand long-term killing dynamics.

The current standard for persister identification at the population level is that the killing curve is at least biphasic, where two time-scales are identified in the time-kill curves [4]. Persisters are identified as the subpopulation with a second, slower killing rate than the rapid death rate of the primary population. If only two timescales are present in the killing dynamics, the population survival time can be extrapolated from the second slow killing rate. Notably, the presence of more than two phases has been demonstrated previously in a few studies [3, 13, 14, 15].. These observations motivate the importance of studying the long-term survival of the antibiotics-tolerant subpopulation, which may not agree with extrapolation from short-term survival. However, most *in vitro* lab research on persisters of fast growing bacteria as *E. coli* is carried out for three to five hours [16, 7, 5], though some studies increase the exposure time to 24–50 hours [2, 15, 17, 3]. Investigation of long-term survival beyond the typical five-hour persister assay might reveal new insights into bacterial killing dynamics that are relevant for the week-long antibiotics treatment of bacterial infections recommended by the WHO [1].

Lastly, the molecular mechanism(s) of persister formation is still unknown [18, 12]. Many intracellular components have been proposed to play a role [19, 16, 20, 21, 22, 23], but so far no single mechanism convincingly explains persister formation. In fact, bacterial persistence presents as a very complex and diverse problem, where the survival fraction could be composed of different subpopulations.

Despite the complexity of persistence, it has been established that stationary phase cultures contain a greater persister fraction than exponentially growing cultures [5]. Stationary phase bacteria may refer to bacteria in a multitude of different physiological states, but is typically associated with starvation stress [24]. Furthermore, the second messenger (p)ppGpp, which accumulates during starvation responses was frequently shown to correlate positively with persistence formation [25, 17, 26, 5, 21]. Hence, it is critical for persistence research to understand the degree to which (p)ppGpp levels affect persistence, and under which circumstances.

The present study investigates persistence in the balanced exponential growth phase where (p)ppGpp levels are relatively low and correlate inversely with the growth rate. It deals with whether *E. coli* forms spontaneous persisters in the exponential phase, their dependence on the growth rate, how long they survive and how their formation relates to (p)ppGpp levels. We followed the long-term survival of *E. coli* K12 populations exposed to a lethal concentration of ciprofloxacin for one week. The growth rate of the *E. coli* population at the time of antibiotics exposure was varied using growth medium with either of two different carbon sources. In addition, a knockout strain was constructed in the wild-type background, removing the gene *relA* and, thus, introducing a (p)ppGpp synthesis deficiency. Furthermore, we compared the killing dynamics with and without a short carbon-starvation period immediately prior to the killing assay. The starvation pulse had a considerable influence on persister formation. This triggered persistence had a very long memory effect on the survival of the population.

## 2. Results

### Long-term persister assay of exponentially growing cells

First, we investigated whether long-term persister cells form during exponential growth in glucose minimal medium. We treated balanced cultures of *E. coli* K-12 with ciprofloxacin and monitored the killing dynamics for one week of antibiotics treatment. Balanced growth was obtained by culturing the cells for more than twenty doubling times in the target medium at 37 °C, keeping the cell density of the culture below an OD_436_ of 0.3 by repeated back-dilutions. Cultures were then treated for a week with 10 μg/mL ciprofloxacin and their killing dynamics were monitored by repeated platings of culture aliquots on antibiotics-free growth medium (See methods for details).

*E. coli* persisters were formed during exponential balanced growth in glucose minimal medium, as seen in Fig. 1. There was a fast initial killing at a rate of about 1/0.3 (h^−1^), with a slower killing rate already after two hours. When we fit a biphasic curve (summation of two exponential functions) to the data up to 7 hours, the second phase of killing is at a rate of about 1/2.3 (h^−1^), shown by a brown line in the inset in Fig. 1. However, for longer times, this fit significantly underestimates the survival time of the bacterial population (Fig. 1). In other words, there is an even slower phase of killing, extending from seven hours to four days. Lastly, from day five to day seven, the remaining cells were killed at a very slow rate, however, this part of the data is less reliable due to the small numbers of recovered colonies.

**Figure 1:**
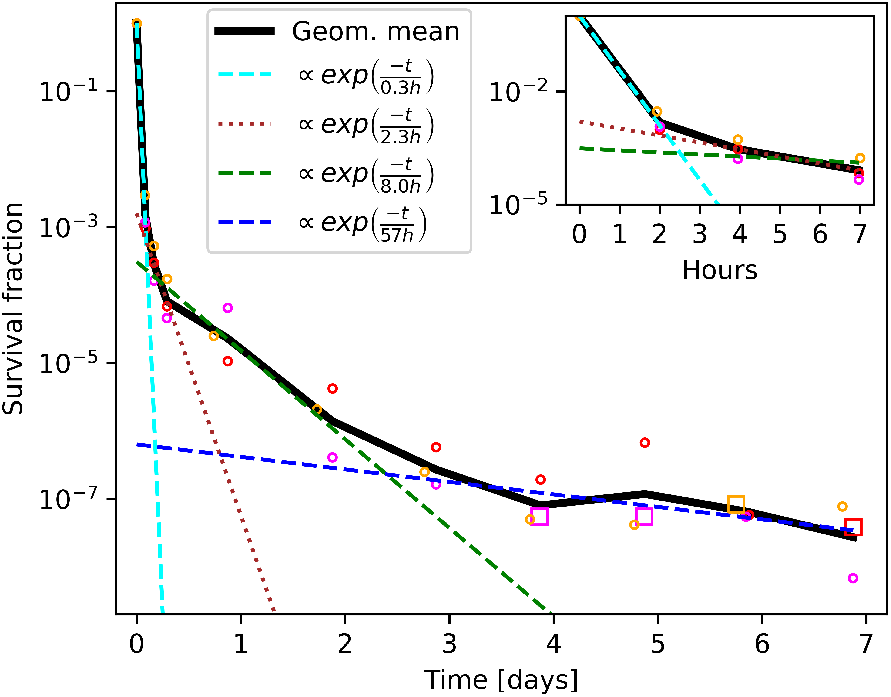
Killing dynamics of exponential phase *E. coli* persisters. Killing dynamics for exponential phase persisters. The bacteria were grown in glucose minimal medium. All three biological replicates are shown for each data point. The black line is the geometrical mean. Each separate phase of a triphasic fit are shown respectively as cyan, green and blue dashed lines. The second phase of a biphasic fit to the first seven hours is shown as a brown dotted line.

To identify the various phases in a quantitative manner, the data was fitted with a sum of exponentials, and the appropriate number of exponentials was chosen with a *χ*^2^ test (see Supplementary text S1 section 2). The double exponential, which corresponds to the biphasic killing dynamics, was rejected as a good fit by the test. Instead, a fit using the triple exponential functions was accepted. The best fit obtained was the first phase of killing at a rate 1/0.3(h^−1^) (a cyan dashed line), the second phase of killing at a rate of 1/8(h^−1^) (a green dashed line), and the third phase of killing at a rate of 1/57(h^−1^) (a blue dashed line). The long-term killing dynamics thus had more than two time-scales, which is only apparent after several days of measurement.

The exponential phase growth rate determines many aspects of bacterial physiology, including the macromolecular composition [8, 27]. The growth rate at the time of antibiotics exposure has previously been linked to short-term survival of antibiotics [10, 11], and could also affect long-term survival. For that reason, the long-term killing assay was repeated with glycerol as the carbon source, which strongly affected the wildtype growth rate. In glucose minimal medium, the wildtype doubling time was 50±1.4 min, while it was 106±3.0 min in glycerol minimal medium. This difference had an impact on the initial phase of killing for up to seven hours, showing slower killing and a higher survivor fraction in glycerol (Fig. 2A inset).

**Figure 2:**
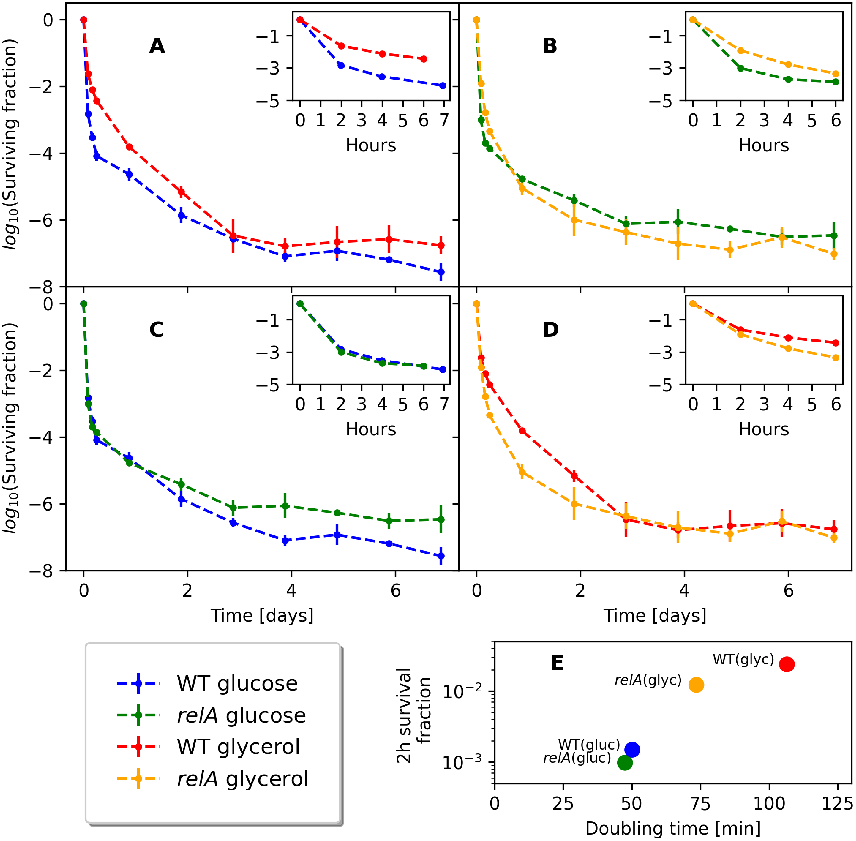
Killing dynamics in different medium with and without the *relA* gene. Each line represents three biological replicates. (A) The wildtype killing dynamics in either glycerol or glucose minimal medium. (B) The Δ*relA* killing dynamics in either glycerol or glucose minimal medium. (C) Killing dynamics in glucose minimal medium for comparison between the wildtype and the Δ*relA* strain. blackThere was no statistically significant difference between the wildtype survivors and the Δ*relA* strain survivors at any of the timepoints (Supplementary text S1 section 2). (D) Killing dynamics in glycerol minimal medium for comparison between the wildtype and the Δ*relA* strain. (E) Survival fraction after two hours of ciprofloxacin treatment compared to the growth rate prior to antibiotics treatment.

However, the long-term killing curve in glycerol minimal medium was not merely a decelerated version of the killing curve in glucose minimal medium. Though significantly more cells survive in glycerol than in glucose for up to a day (Fig. 2A), the survival curve in glycerol medium had a steeper slope than in glucose medium, resulting in comparable surviving fractions after 2 to 3 days. In fact, in two of the three biological replicates of wildtype cultures grown in glycerol, almost no survivors were observed after 3 days of killing (see Supplementary text S1 section 1), supporting further that the survival in glycerol minimal medium is not more than that in glucose minimal medium after 3 days.

Overall, the wildtype killing dynamics in glycerol had more than two time scales, with a best fit of four separate phases of killing (Supplementary text S1 section 2).

### Deletion of relA affects the killing dynamics

In glycerol minimal medium, the steady state (p)ppGpp level of the wildtype strain is higher than in glucose minimal medium [28]. Because the (p)ppGpp level has been frequently associated with persister formation, we next aimed at investigating the effect of the initial (p)ppGpp level on long-term survival. *E. coli* encodes the primary (p)ppGpp synthetase RelA and the secondary (p)ppGpp synthetase SpoT, the latter of which is bifunctional as a (p)ppGpp hydrolase. The nutrient-dependent steady-state growth rates are inversely related to the concentrations of (p)ppGpp, both for *relA*^+^ an *relA*^−^ strains [28], and for many carbon sources, the growth rates and (p)ppGpp levels of *relA*^+^/*relA*^−^ strain pairs are indistinguishable due to (p)ppGpp synthesis by SpoT. However, in low energy carbon sources like glycerol or acetate, SpoT produces insufficient (p)ppGpp to reduce the growth rate when RelA is missing, leading to an enhanced growth rate of the RelA mutant strain relative to the wildtype [28]. We constructed a Δ*relA* mutant to clarify the role of (p)ppGpp in the killing dynamics.

As expected, the difference between the growth rate in glucose and in glycerol minimal medium was smaller for the Δ*relA* strain than the wildtype, with a doubling time of 47±1.5 min in glucose minimal medium and 74±1.6 min in glycerol minimal medium. The survival after 2 hours is positively correlated with the doubling time (Fig. 2E), consistent with the previous observations that the initial killing rate decreases with the doubling time [11]. The short term (up to 4 hours) survival under antibiotics exposure was also correlated with the growth rate in the Δ*relA* strain (Fig. 2 inset).

Like wildtype cells, the Δ*relA* mutant formed long-term survivors in both glucose and glycerol minimal medium with more than two phases of killing (Fig. 2B, see Supplementary text S1 section 2 for statistical analysis). In glucose, the two strains grew at similar rates, and the early killing dynamics of the Δ*relA* mutant were very similar to that of the wildtype (Fig. 2C). In glycerol, the faster-growing Δ*relA* mutant showed a significantly lower level of persisters than the wildtype in the initial phase of killing up to one day (Fig. 2D, see Supplementary text S1 section 2), indicating the importance of growth rate, or ppGpp level, for persister formation in this phase. Interestingly, the long-term survival of the Δ*relA* mutant and wildtype were comparable at later times (after 3 days) in glycerol medium, and somewhat higher than the wildtype strain in glucose medium. Thus, survival in the long term is not simply dependent on the population growth rate at the time of antibiotics exposure.

### A starvation pulse prior to the antibiotic application affects the long-term persistence of wildtype cells in glucose minimal medium

A sudden downshift of the carbon source is known to give a spike of the (p)ppGpp level in the wildtype strain just after the downshift, while the spike is significantly lower in a RelA strain [29]. We then wondered if a short pulse of carbon source starvation to the exponentially growing cells prior to the killing assay would give a quantifiable difference in the long-term persistence between the wildtype strain and the Δ*relA* strain. If the starvation pulse increases the persisters, it would be considered as triggered persistence, and the current study would allow us to quantify how long such triggered persisters last.

In order to test this, part of the cultures in balanced growth were filtered into growth medium without a carbon source and starved for 1 hour, before the carbon source was replenished (Figure 3A). Figure 3B shows that the 1-hour starvation pulse resulted in a quick rise of the ppGpp level peaking at about 15 min after the downshift for the wildtype strain in glucose medium, while only a mild increase of the ppGpp level was seen in the Δ*relA* strain. Antibiotics were added simultaneously with the carbon source replenishment (Fig. 3A, see methods for details). Remarkably, the short carbon starvation prior to the addition of antibiotics had long-term effects on the killing dynamics. This was especially visible for the wildtype strain grown in glucose minimal medium, where the brief starvation period reproducibly resulted in almost 100-fold more persisters for up to four days. The difference was abolished by removing *relA*, as seen in Fig. 4B; the Δ*relA* strain only exhibited increased survival for the first six hours following starvation. As such, the long-term memory of the starvation pulse is seemingly a *relA*-dependent effect. However, the Δ*relA* strain in the steady state growth in glucose had more long-term survivors than the wild-type strain, and the downshift brought the wildtype strain survival fraction to a level similar to the Δ*relA* strain.

**Figure 3:**
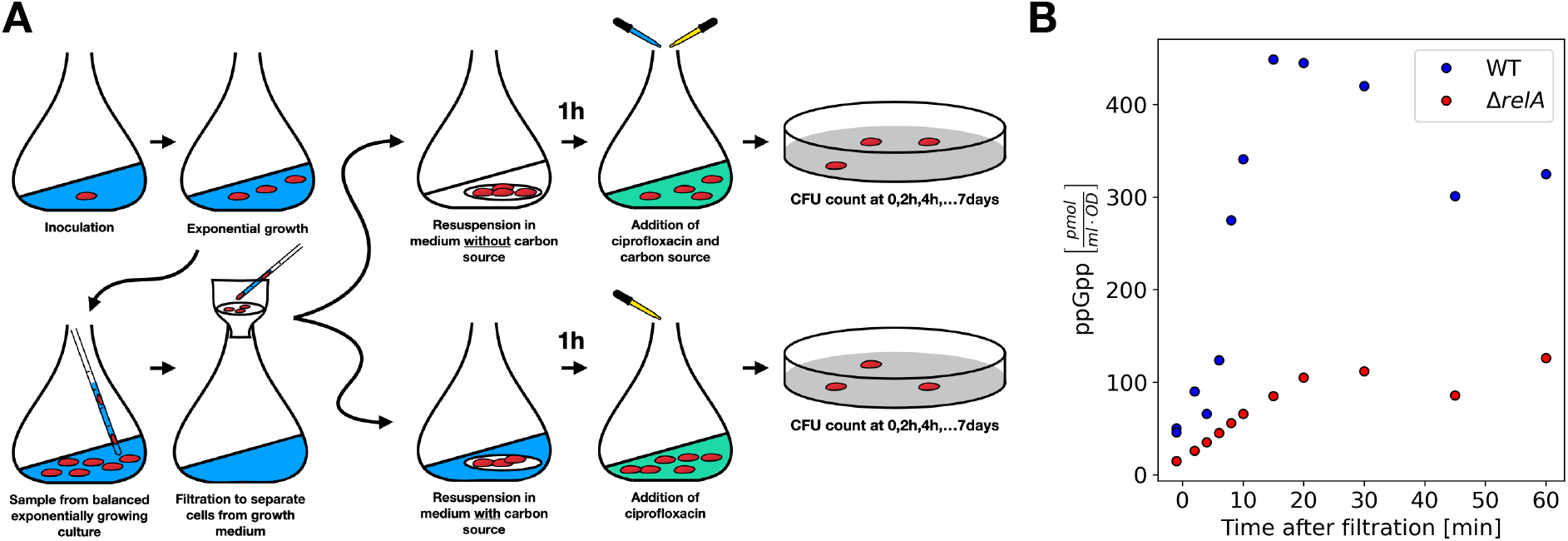
The experimental setup for the long-term persistence assay. (A) Cells were in balanced exponential growth prior to the killing assay. A part of the culture was filtered and then resuspended in medium with or without the carbon source, and incubated for 1 hour. After 1 hour, the ciprofloxacin was added to both of the samples, and at the same time the carbon source was added to the the culture that has been starved fro 1 hour. The first sample was taken just before the additions. After that, samples were taken at 2,4,6/7, 21 +24·*n* hours after the antibiotic addition for *n* ∈ [0; 6]. The samples were washed and plated on agar plates containing the target medium. (B) The 1-hour time course of ppGpp level for the culture grown in the glucose medium, filtered, and resuspended in fresh medium without carbon source. The time zero is the time of the resuspension. The wild type (blue circles) shows clear peak around 15 minutes after the resuspension. ΔrelA strain shows only mild increase of the ppGpp level. The phosphoImager scan of TLC plates used to quantify ppGpp levels is shown in Supplementary Fig. S7.

**Figure 4:**
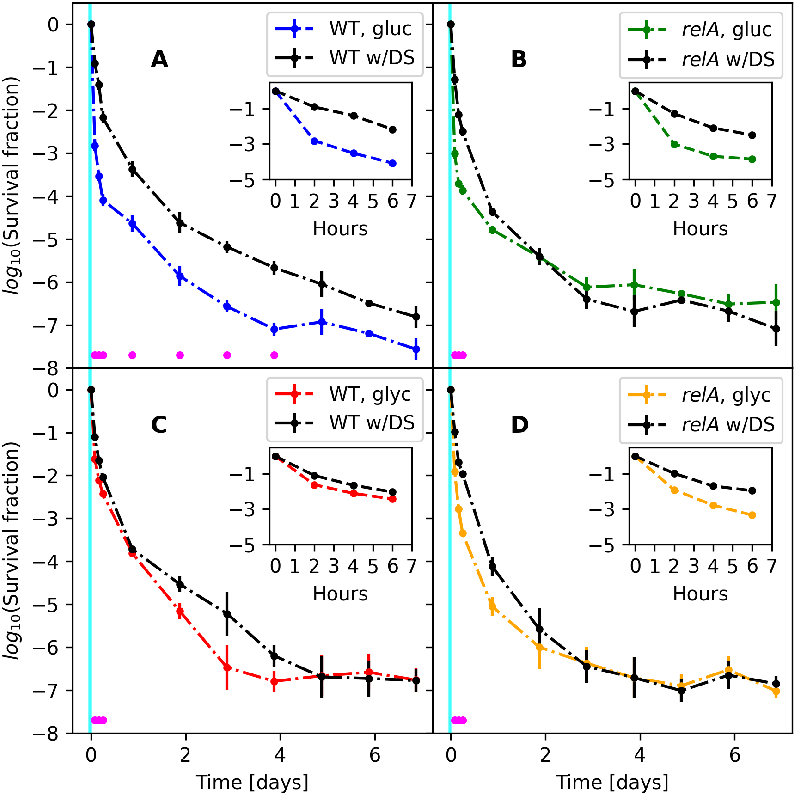
The effect of a short starvation-pulse on the survival dynamics. All strains and conditions are shown with and without the downshift. The black line is with the downshift, the colored line is without the downshift. The magenta dots represent statistically different datapoints as tested by an unequal variances two-sided t-test. The cyan interval illustrates the downshift period prior to the killing assay. (A) The wildtype in glucose. A downshift in glucose significantly enhances wildtype survival for four days and seems to be increased for up to all seven days. (B) The Δ*relA* mutant in glucose. (C) The wildtype in glycerol. (D) The Δ*relA* mutant in glycerol.

The effect of the downshift was smaller in the glycerol medium (Fig. 4CD). In the wildtype strain, the average persister level with downshift was higher up to 4 days, but the statistical significance of the difference was confirmed only up to 7 hours due to the larger data scatter for later time points (Fig. 4C). The effect of downshift disappeared faster in the Δ*relA* strain already after 2 days (Fig. 4D).

## 3. Discussion

We expanded the understanding of bacterial killing dynamics with a long-term persister assay. The use of minimal medium facilitated the formation of long-term persisters during exponential growth, in contrast to growth and killing in rich medium [7]. Spontaneous persisters were observed during the exponential growth phase, both in glucose and glycerol minimal medium, and in some cases they survived at least one week. This long-term survival does not require *relA*, although the residual (p)ppGpp synthesized by SpoT is likely necessary. In fact, there is an increase in long-term survival of the Δ*relA* mutant in glucose.

We have shown that a one-hour starvation pulse prior to the addition of the antibiotic affects long-term survival. The finding that a short starvation pulse gives a long-term effect is consistent with a previous study of a temporal nitrogen downshift prior to antibiotics treatment, which was shown to elevate the persister level at 24 hours in a *relA* dependent manner [30]. Also, a few other studies have previously shown that triggered persistence can be *relA* dependent [25, 21]. Our study demonstrated that the memory can be remarkably long-lasting, as one-hour carbon starvation gave an increase in survival for at least 4 days in the wildtype strain grown in glucose medium. The molecular mechanism underlying this long-term memory is yet to be investigated, but in all likelihood it is linked to the abrupt RelA-mediated rise in (p)ppGpp upon starvation, since the starvation-pulse effect on long-term survival was abolished in the Δ*relA* mutant. In further support of this hypothesis, there was no long-term effect when glycerol was used as the carbon source, which could be due to the high basal level of (p)ppGpp in glycerol relative to glucose minimal medium [8]. The sensitivity of the survival fraction to the rather short perturbation may be consistent with the idea that there is a threshold in some molecule concentration to determine if the cell becomes a persister or not [31, 32], since a small perturbation can have a major impact on the occurrence of rare expression patterns that exceed an extreme threshold [33]. This observation also alerts us that blacka small perturbation in the experimental procedure may strongly affect the result of persister assays.

This study shows that the long-term killing of *E. coli* in ciprofloxacin is not adequately described by biphasic dynamics. At least three phases of killing were present in the data. Thus, despite the emphasis on at least a biphasic behaviour to define persistence [4], a third, or even fourth, phase of killing occurs that may even be more clinically relevant. The presence of additional phases also means that the population survival time will be underestimated by predictions from the biphasic killing assumption. Indeed, it is not sufficient to measure killing dynamics for only five hours and then extrapolate the population survival time from there.

The detailed molecular mechanism of the observed persistence is not the focus of the current study. Nevertheless, it is worth mentioning that ciprofloxacin has been reported to induce persistence via toxin activation through the SOS response in the killing dynamics up to 6 hours [34]. The existence of more than two killing phases indicates that different mechanisms may play roles for longer-term persistence on top of the previously studied ones. For the future study of the molecular mechanisms of persistence, attention should be paid to which time scale of the survival the pathway affect.

The population growth rate was found to be positively correlated with the killing rate in the initial phase. However, the correlation was diminished in the longer term survival, and lost in the third phase of killing. In shorter persister assays, a difference in growth rate, such as between different mutants, might strongly confound results when analyzing persistence fractions. These differences seem smaller and less relevant in later phases of killing.

This investigation of long-term killing dynamics has added to the concept of bacterial persistence as a complex phenomenon. During long-term antibiotics treatment, different mechanisms could account for bacterial survival on different timescales (an hour, a day, a week), although the (p)ppGpp level at the time of initial exposure to the antibiotic seems to be important in all cases. As such, persistence seems to be a time-dependent phenomenon, where different survival mechanisms account for different bacterial life spans. The presence of several phases in the killing dynamics begs the question if a more extended concept should replace the simplified concept of bacterial persistence.

## 4. Methods and Materials

### Strains

The strain MAS1081 (MG1655 *rph*^+^ *gatC*^+^ *glpR*^+^) was used as the wildtype [35]. The Δ*relA*, MAS1191, is MAS1081 made Δ*relA*251 :: *Kan* by P1 transduction from CF1651 [36] followed by selection on kanamycin.

### Long-term killing assay

A single colony of the *E. coli* strain grown on an agar plate was incubated overnight in the target medium (MOPS minimal medium with either glucose or glycerol as the carbon source [37]. See the Supplementary text S1 section 1 for the recipe). This was done for each biological replicate. The overnight culture was diluted 1 : 10^7^ in 10 mL target medium in a 100 mL Erlenmeyer flask. The flask was continuously shaken at 160 RPM in a 37°C room. Hours later, the culture was diluted further, at least 1 : 10^2^, in 100 mL preheated medium in a 1L Erlenmeyer flask, reaching a total dilution of at least 1 : 10^9^. Once the OD_436_ reached a detectable level of 0.01, at least 5 samples were measured at different timepoints to establish a growth rate (supplementary Fig. S4). The OD_436_ was consistently kept below 0.3. A part of the culture, typically around 20 mL, was filtered, and the cells on the filter were resuspended in 40 mL of the target medium (w/o carbon source) in a 300 mL Erlenmeyer flask. As such, both the control and the starved culture were exposed to filtration and resuspension. The medium volume never exceeded 14 percent of the flask volume. The starvation was verified by measuring OD_436_, to confirm either increase in biomass (control) or no growth (starving culture). See the Supplementary text S1 section 1 for confirmation of the downshift. After one hour of starvation, a sample was taken immediately before ciprofloxacin (10 *μ*g/mL) was added along with the carbon source, that was replenished to end the downshift. Samples were taken at times 2,4,6/7, 21+24·*n* hours for *n* ∈ [0; 6]. The samples were put on ice for a few minutes and then centrifuged for ten minutes at 4°C at 10,000*g*. The supernatant was removed and the cell pellet was resuspended in room temperature MOPS buffer with no supplements. The sample was diluted appropriately, never more than 1 : 100 per step, corresponding to 10 *μ*L in 990 *μ*L. The sample was plated with 200 *μ*L per plate on minimal medium plates containing the target medium. The plates were kept at 37°C for at least one week and all colonies were counted. The whole experiment is illustrated in figure 3. Detection limit of each experiment is presented in supplementary Fig. S6.

### Controls

After each completed experiment (7 days), 200 *μ*L of culture, still containing antibiotics, was spread on a plate with the target medium. This was left at 37°C for at least 7 days, and no growth confirmed the absence of a growing resistant culture in the flask. In addition, the activity of the antibiotics in the culture was tested after seven days by dropping 20 *μ*L on a lawn of growing *E. coli*.

### Analysis of time-scales in the killing dynamics

Time-scales in the killing dynamics were statistically identified by fitting a sum of exponential functions to the data. The model with the least number of parameters, that could not significantly be rejected, was then chosen [38]. The functional form of the models was a sum of exponential functions. The number of exponential functions in the sum corresponded to the number of time-scales. A biphasic killing curve would for example be well fitted by the sum of two exponentials. The *χ*^2^-test for goodness of fit is used for identifying the appropriate model using the Minuit minimization software [39, 40] (see the Supplementary text S1 section 2 for specifics of the analysis).

### Statistical analysis

All killing curves are based on three biological replicates. The mean is determined as the mean of the logarithmic values, which corresponds to the geometric mean. This is done to get a more adequate mean-value representation in log-space. The uncertainties are also calculated as the standard deviations of the log transformed values. Whenever a datapoint had the value zero, which happened frequently, that value was replaced with the detection limit, to get a sensible value in log-space. An unequal variance two-sided t-test was used to determine significant differences between two datapoints at the same timepoint (*P <* 0.05).

### ppGpp measurements

The measurements Were performed essentially as described in [41] and used in [42, 43, 44]. In short: Cultures were grown for two generations in the presence of 75*μ*Ci/mL 32P-phosphate at a total phosphate concentration of 0.33 mM. At the time of starvation, cultures were filtered, washed in medium without glucose and phosphate and resuspended in medium without glucose but containing 32P at the same specific activity as during growth. These steps were performed at 37°C and lasted less than 2 min. For determination of the nucleotide pools, 100 *μ*L of culture was harvested into 20*μ*L 2M formic acid at 0°C. After centrifugation the nucleotides in the supernatant were separated by chromatography on polyethyleneimine-cellulose plates. The activities of the individual spots were quantified by PhosphoImager scans (Typhoon Phosphor Imager FLA7000 (GE Healthcare)) of the plates. The specific activity of the signal was determined from a medium sample from the individual cultures spotted onto the same brand of plates.

## Acknowledgments

MSS and NM thank S. Semsey for fruitful discussions. This work was supported by the Danish National Research Foundation (DNRF120), the Independent Research Fund Denmark (8049-00071B and 8021-00280B), and the Villum foundation (00028054).

