## Supplementary material for "Existence of log-phase Escherichia coli persisters and lasting memory of a starvation pulse"

**3 ppGpp level during glucose downshift w/o *relA*** **15**

### 1 Details of the experiments

#### 1.1 Growth medium

##### 1.1.1 Mops minimal medium

- 100 mL 10X MOPS, which contains the following concentrations according to [4].
  - 40 mM MOPS
  - 4 mM tricine
  - 1.32 mM $K_2HPO_4$
  - 9.52 mM $NH_4Cl$
  - 0.523 mM $MgCl_2$
  - 0.276 mM $K_2SO_4$
  - 0.01 mM $FeSO_4$
  - 0.5 $\mu$M $CaCl_2$
  - 50 mM NaCl
  - trace metals (see [5])
- Carbon source (one of either)
  1. 10 mL 20% glucose (for 0.2 % w/v glucose minimal medium)
  2. 8 mL 50% glycerol (for 0.4 % v/v glycerol minimal medium). The 50% glycerol stock solution is stored at -20 °C.
- Fill up to 1L milliQ water

##### 1.1.2 Mops minimal medium plates

- Add 15 g Bacto agar (autoclaved with the milliQ water) to the Mops minimal medium recipe.

#### 1.2 Each experiment shown separately

Each biological replicate of the glucose experiments are shown in Fig. S1, whereas the replicates of glycerol experiments are shown in Fig. S2. If the CFU was zero, corresponding to no colonies counted, the value zero was replaced by the detection limit.

| Experiment | Carbon source | Strain | Doubling time (min) |
|---|---|---|---|
| A | Glucose | WT | 46.8 |
| B | Glucose | WT | 51.4 |
| C | Glucose | WT | 52.4 |
| D | Glucose | $\Delta relA$ | 47.8 |
| E | Glucose | $\Delta relA$ | 44.3 |
| F | Glucose | $\Delta relA$ | 50.6 |
| G | Glycerol | WT | 103 |
| H | Glycerol | WT | 102 |
| I | Glycerol | WT | 114 |
| J | Glycerol | $\Delta relA$ | 69.8 |
| K | Glycerol | $\Delta relA$ | 74.3 |
| L | Glycerol | $\Delta relA$ | 76.5 |

Supplementary Table S1: Metadata for all long-term killing experiments

#### 1.3 Metadata for each experiment

The metadata for all experiments, providing results for this study, are presented in table S1. The doubling time reported in table S1 is from the exponential growth, prior to the downshift and the actual experiment. The doubling time is thus the same for both the experiments with and without a carbon source downshift.

Following the balanced exponential growth, a carbon source downshift was induced for one hour, which is compared to a control culture, that did not experience a downshift. The downshifts in the experiments were verified by OD measurements. The idea was to verify growth in the control cultures, with the carbon source present, and no growth in the cultures without the carbon source present. The OD was measured twice during the one hour downshift, to confirm either an increase or no change in biomass. The absorbance measurements verified that a starvation took place for all the experiments, that were supposed to experience a downshift. It also verified that growth, with a doubling time approximately close to the expected value, took place in the control cultures.

#### 1.4 Growth curves for all cultures

[h!]

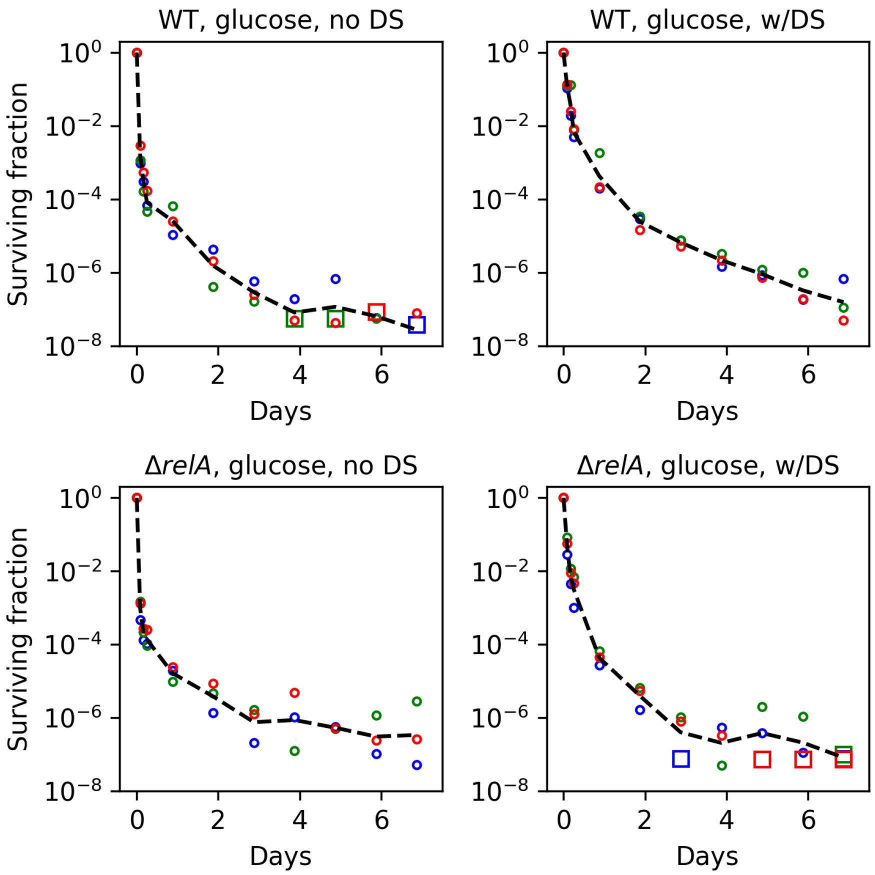

Supplementary Fig. S1: Each replicate shown for all experiments in glucose. The black line represents the geometric mean. Each datapoint represents a biological replicate. The circles are datapoints with a nonzero CFU count. The squares represent a CFU count of zero, where the zero is replaced by the detection limit. Data from experiments with a sugar downshift are labeled ”w/DS”, whereas ”no DS” refers to experiments without a downshift.

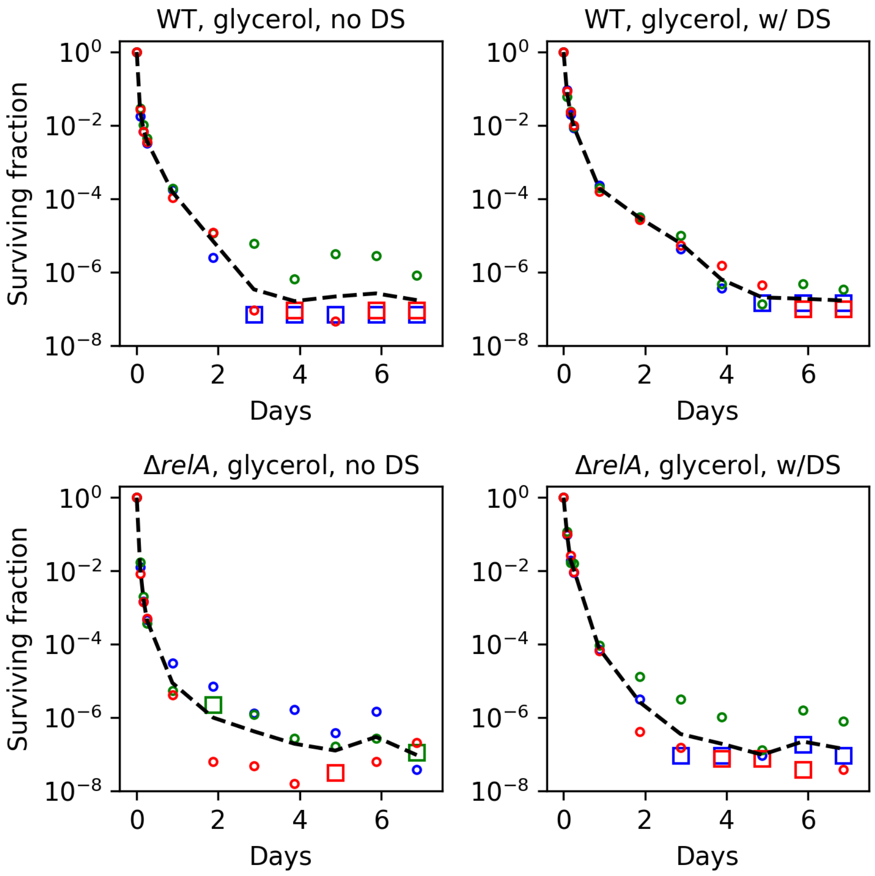

Supplementary Fig. S2: Each replicate shown for all experiments in glycerol. The black line represents the geometric mean. Each datapoint represents a biological replicate. The circles are datapoints with a nonzero CFU count. The zero represent a CFU count of zero, where the zero is replaced by the detection limit.

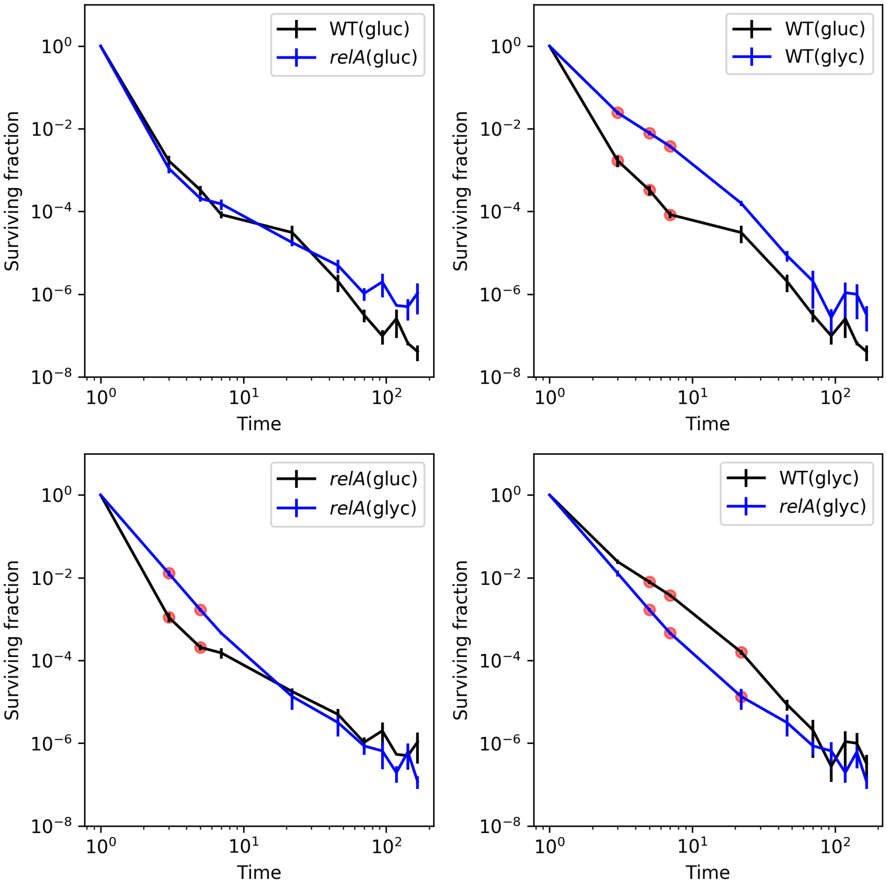

Supplementary Fig. S3: Comparison of killing dynamics in exponential phase persisters between the two different strains and two different conditions. The fourth datapoint of the wildtype in glucose minimal medium is corrected in time, since the sampling time was 7 hours, compared to 6 hours for the others. A unequal variance t-test was done between each sampling. Only the initial killing was significantly different for some of the conditions. The time-axis starts at 1 hour to show the data in a log-log representation.

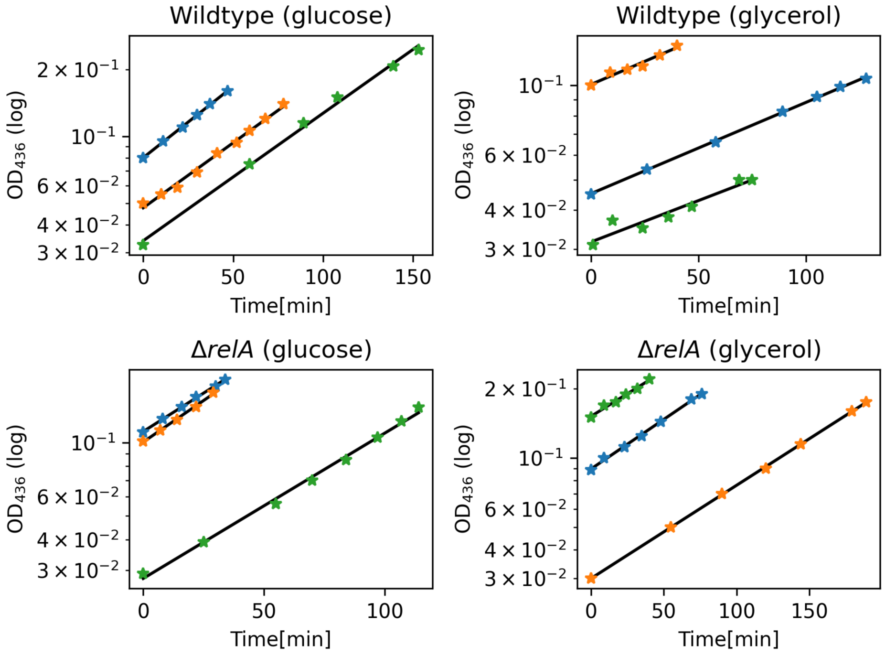

Supplementary Fig. S4: Growth curves for all cultures used for experiments in this study. Two subcultures were investigated from each of these cultures: one with a downshift and one without.

### 2 Data analysis

#### 2.1 Analysis of time-scales in the killing dynamics

Time-scales in the killing dynamics are statistically identified by fitting a sum of exponential functions to the data. The model with the least number of parameters, that cannot significantly be rejected, is then chosen, following [6]. The functional form of the models are as follows:

$$
\begin{aligned}
Model1(t, a_0) &= \exp(-a_0 \cdot t) \\
Model2(t, a_0, a_1, b_1) &= (1 - b_1) \cdot \exp(-a_0 \cdot t) \\
&\quad + b_1 \cdot \exp(-a_1 \cdot t) \\
Model3(t, a_0, a_1, a_2, b_1, b_2) &= (1 - b_1 - b_2) \cdot \exp(-a_0 \cdot t) \\
&\quad + b_1 \cdot \exp(-a_1 \cdot t) \\
&\quad + b_2 \cdot \exp(-a_2 \cdot t) \\
Model4(t, a_0, a_1, a_2, a_3, b_1, b_2, b_3) &= (1 - b_1 - b_2 - b_3) \cdot \exp(-a_0 \cdot t) \\
&\quad + b_1 \cdot \exp(-a_1 \cdot t) \\
&\quad + b_2 \cdot \exp(-a_2 \cdot t) \\
&\quad + b_3 \cdot \exp(-a_3 \cdot t)
\end{aligned}
$$

Each exponential function contains a specific time-scale which corresponds to each exponent. A biphasic killing curve would for example be well fitted by the sum of two exponentials. To statistically determine which model fits to the data, each model is fitted to the data by using least squares and the Minuit optimization software [1,2]. The $\chi^2$ is then calculated for each of these models. The degrees of freedom correspond to the number of datapoints with the subtraction of the number of parameters for fitting and one degree of freedom for normalizing the data. Thus, the probability that the data does not correspond to the model is calculated using the $\chi^2$ cumulative distribution. The p-value was chosen to be 0.05. For the wildtype strain grown in glycerol, the Model4 was an appropriate fit, for the three other datasets, the Model3 was an appropriate fit. The parameters estimated are given in table S2. The fits are shown in figure S5.

| Parameters | $a_0$ $[1/h]$ | $a_1$ $[1/h]$ | $a_2$ $[1/h]$ | $a_3$ $[1/h]$ | $b_1$ | $b_2$ | $b_3$ |
|---|---|---|---|---|---|---|---|
| Model3, WT, gluc. | 3.30 | 0.123 | 0.0117 | - | 3.32e-4 | 3.22e-7 | - |
| Model4, WT, glyc. | 2.30 | 0.301 | 0.136 | 1.59e-3 | 0.0211 | 2.48e-3 | 2.41e-7 |
| Model3, *relA*, glucose | 3.59 | 0.129 | 0.0155 | - | 2.88e-4 | 4.00e-6 | - |
| Model3, *relA*, glycerol | 2.29 | 0.302 | 0.0157 | - | 4.26e-3 | 1.35e-6 | - |

Supplementary Table S2: Parameters estimated by MLE using Minuit.

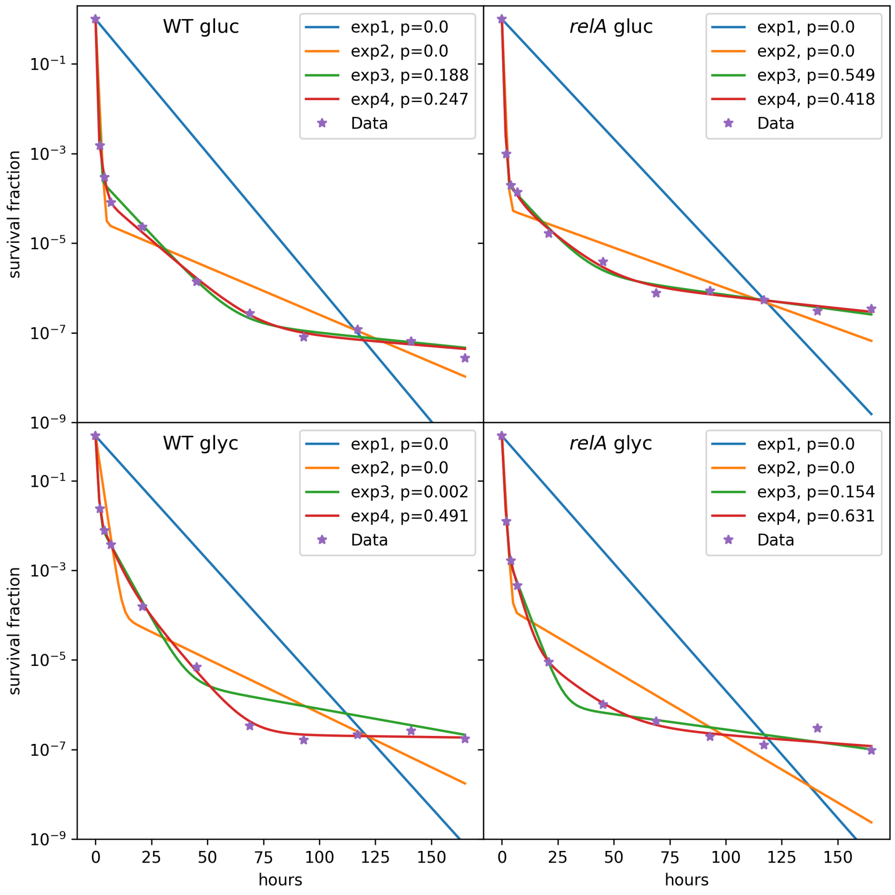

Supplementary Fig. S5: The various exponential models fitted to the data.

##### 2.1.1 Display of fits in figure 1 in the main text

The functions shown in figure 1 in the main text all stem from the fit of Model3 to the data. Model3 is given by equation 1.

$$Model3 = (1 - b_1 - b_2) \exp(-a_0 t) + b_1 \exp(-a_1 t) + b_2 \exp(-a_2 t) \tag{1}$$

The functions displayed are then given by the three terms of the fit shown separately, as given by equation 2,3 and 4.

$$term1 = (1 - b_1 - b_2)\exp(-a_0 t) \tag{2}$$

$$term2 = b_1 \exp(-a_1 t) \tag{3}$$

$$term3 = b_2 \exp(-a_2 t) \tag{4}$$

This is to illustrate the contribution from each exponential time-scale. In addition, the second phase of a biphasic fit to the data shown in the inset, is also shown in the main figure and in the inset.

##### 2.2 Correction for difference in sampling times

The majority of samples were obtained at approximately hours $2, 4, 6, 21 + 24 \cdot n$ where $n \in [0; 6]$. The wildtype glucose experiments were sampled at approximately $2, 4, 7, 21 + 24 \cdot n$, but one of the experiments in glucose had the following timepoints $2, 4, 7, 18 + 24 \cdot n$ where $n \in [0; 6]$. We corrected this dataset, by calculating a straight line between the datapoint of interest and the following datapoint in log space. We used this straight line to extrapolate the value to the timepoint of interest. All statistical testing is done with and without this correction, to make sure it did not affect the results.

| Experiment | $T_{original}$ | $T_{corrected}$ | $\log_{10}$(surv. frac.)$_{ori}$ | $\log_{10}$(surv. frac.)$_{cor}$ |
|---|---|---|---|---|
| A | 6.96 | 6 | -4.166 | -3.961 |
| B | 6.93 | 6 | -4.337 | -4.162 |
| C | 7 | 6 | -3.764 | -3.604 |
| C | 17.8 | 21 | -4.602 | -4.750 |
| C | 41.5 | 45 | -5.681 | -5.811 |
| C | 66.3 | 69 | -6.602 | -6.680 |
| C | 90.6 | 93 | -7.301 | -7.309 |
| C | 114.6 | 117 | -7.380 | -7.349 |
| C | 138 | 141 | -7.079 | -7.082 |

Supplementary Table S3: Data corrected due to deviating sampling protocol. The corrected values are used for calculation of statistical values and to perform test for differences between datapoints of the various conditions. The last datapoint in experiment C is not corrected, since there was no following datapoint to use for the correction, but here it is assumed that little change happened during three hours, compared to the seven days of killing.

### Each experiment with detection limit

WT, glycerol, w/ DS

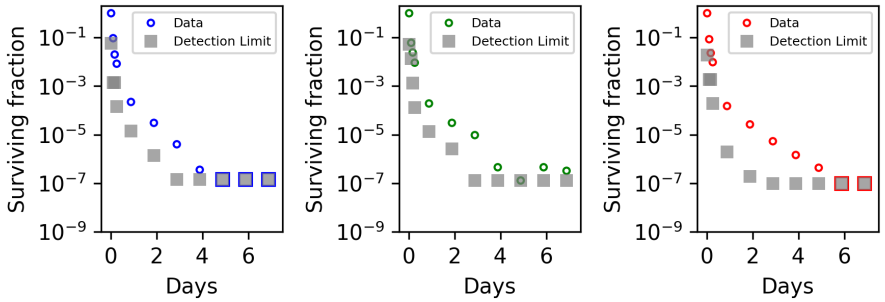

WT, glycerol, no DS

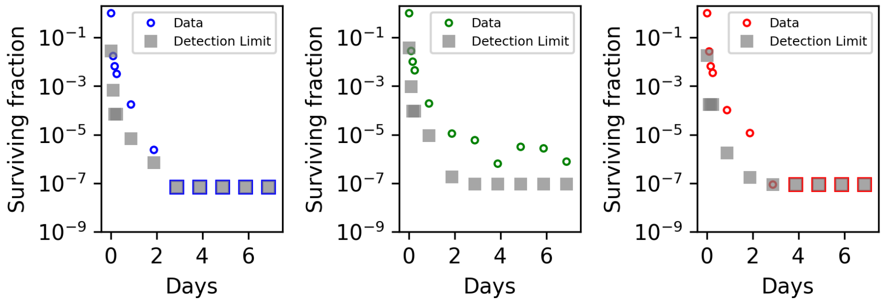

$\Delta$*relA*, glycerol, w/DS

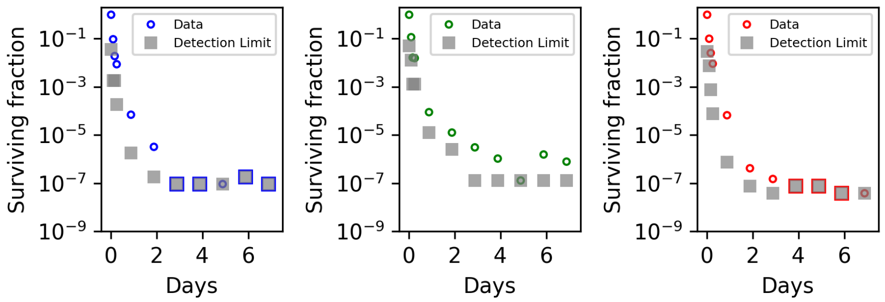

$\Delta$*relA*, glycerol, no DS

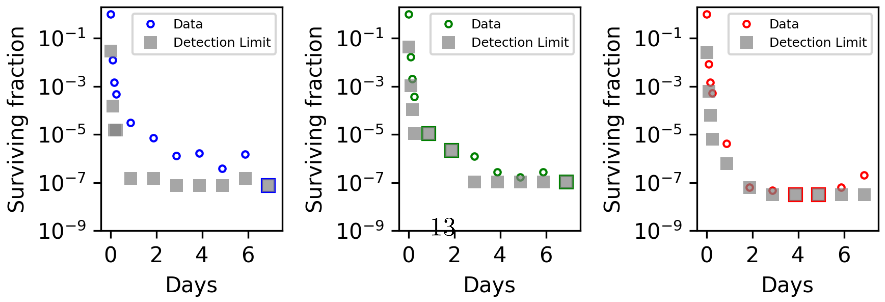

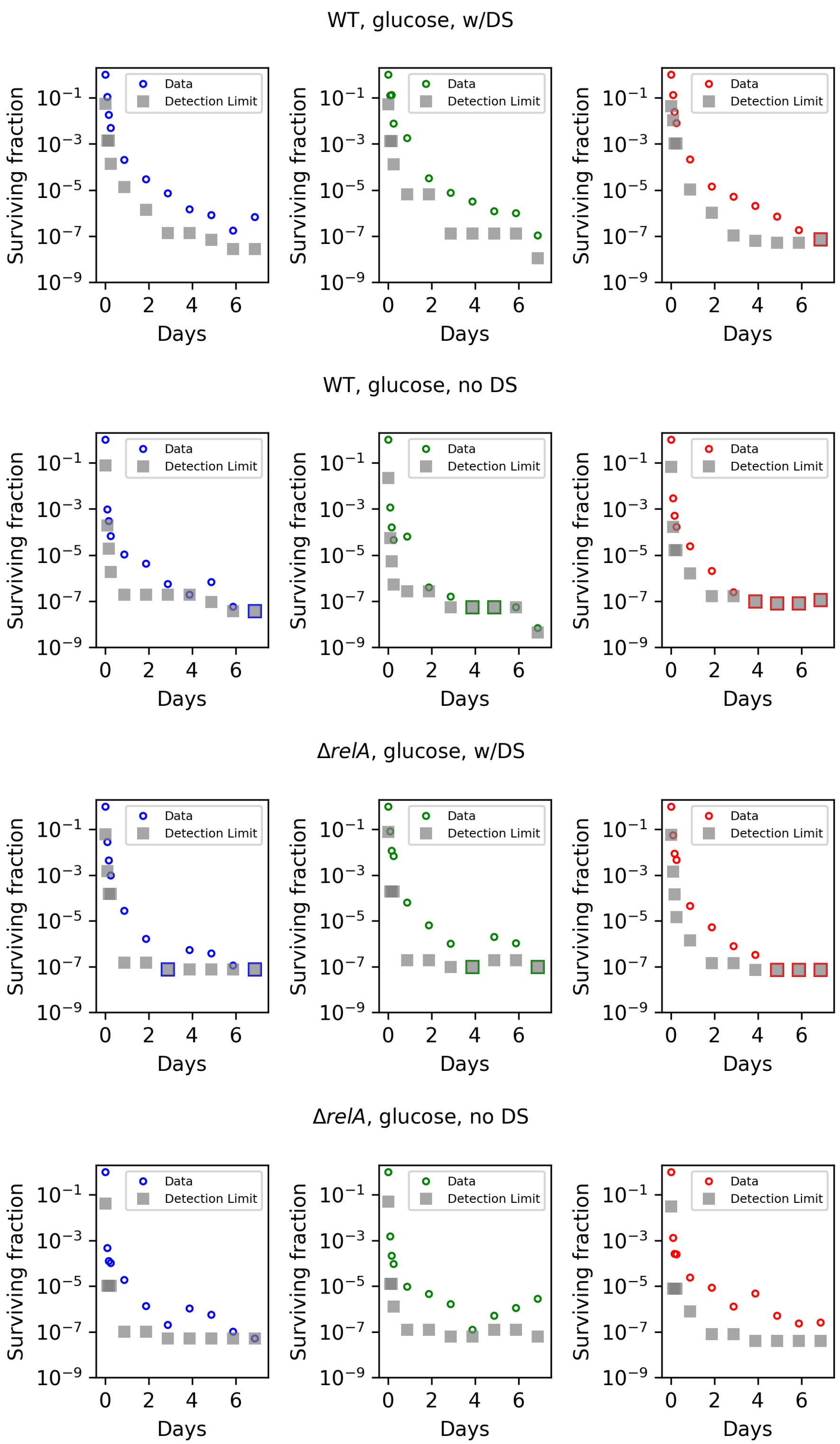

Supplementary Fig. S6: Data of each experiment with detection limit.

#### 3 ppGpp level during glucose downshift w/o *relA*

It was previously shown that a glucose downshift induces a spike in the ppGpp level [3]. However, this effect was to some extent dependent on *relA*. Lazzarini et al. showed that a wildtype strain would form a significantly higher peak during a glucose downshift than the corresponding $\Delta relA$ strain. We wanted to confirm this for our strains and experimental setup. The measurements of ppGpp during a glucose downshift yielded dynamics comparable to those obtained by Lazzarini et al. [3]. The intracellular level of ppGpp reached a higher peak in our wildtype strain than in our $\Delta relA$ strain during a one hour glucose downshift as seen in figure S7.

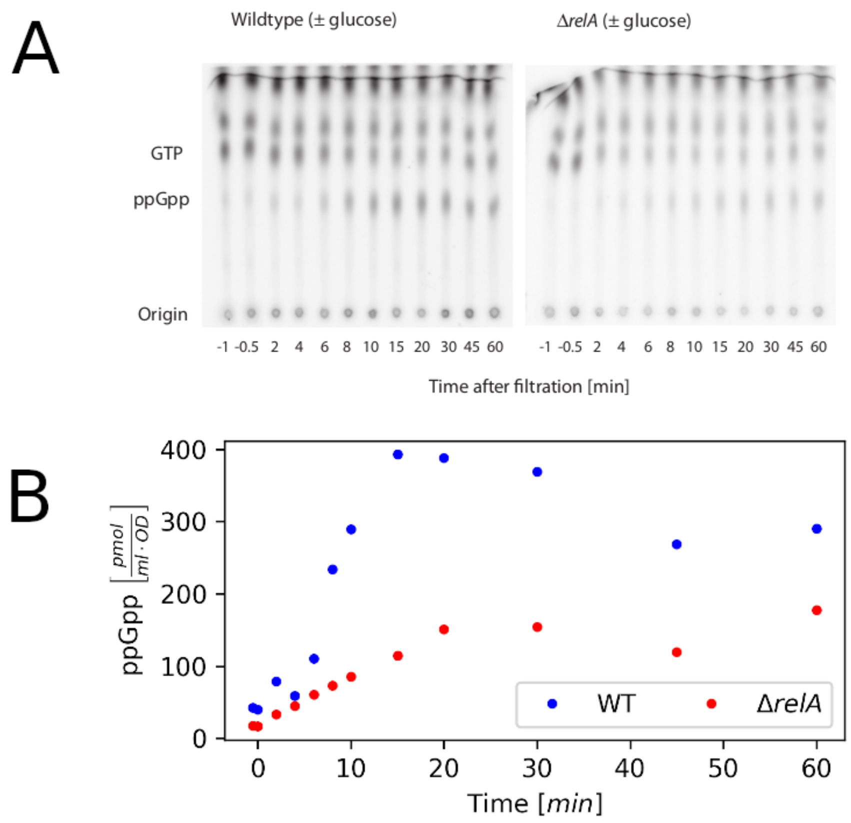

Supplementary Fig. S7: PhosphoImager scan of TLC plates used to separate nucleotide pools in samples from strains in glucose downshifts. The radioactivity in the individual ppGpp spots were quantified to estimate the ppGpp content of the cultures (shown in panel B)
